# Metagenome-assembled genomes (MAGs) revealed a functionally stratified microbiome in Jeevamrit, enabling co-operative nutrient cycling and rhizospheric growth promotion in the natural farming practices

**DOI:** 10.64898/2026.04.21.719941

**Authors:** Draksha Agwan, Ayush G Jain, Anita Surendra Singh, Balaram Mohapatra, Jagat Rathod

## Abstract

Jeevamrit, a fermented liquid microbial bioinoculant, is increasingly recognized as a soil and plant growth enhancer in sustainable agricultural practices such as zero-budget natural farming; however, the genetic pool attributed to the functionality by microbial constituents remained poorly resolved. In this study, we reconstructed 16 high-quality metagenome-assembled genomes (MAGs) from Jeevamrit under two critical mixing regimes to elucidate the contributions of key taxa affiliated with Pseudomonadota, Bacillota, and Bacteroidota to nutrient cycling and plant growth promotion. Functional annotation revealed a stratified (upper-middle-lower) metabolic organization with interdependent interactions driving combinatorial functionality. Upper-layer MAGs, including *Klebsiella* and *Pseudaeromonas* exhibited organic polymer degradation, glycolytic and oxidative carbon metabolism, respiratory versatility with nutrient acquisition traits such as nitrogen fixation and phosphate/iron solubilization. In middle and lower-layer, *Trichococcus, Clostridium*, and *Veillonella* displayed fermentative and reductive metabolisms that facilitate the turnover of partially degraded organic matter and production of organic acids, nitrogen transformations, and metabolic cross-feeding under fluctuating redox conditions. Phylogenetic and taxono-genomic analyses support the designation of eight MAGs as novel species (sp. nov.), for which new names are proposed. A consensus genetic map deciphered traits linked to phytohormone biosynthesis (IAA, cytokinins), quorum-sensing-mediated rhizosphere colonization, and abiotic stress tolerance. Ultimately, this culture-independent metagenome study underpins field-relevant mechanistic insights into an indigenous microbial inoculant, highlighting its potential as a locally adapted solution for sustainable agriculture.

**Importance:** Microbial bioinoculants such as Jeevamrit are increasingly used in sustainable agriculture, yet their functional basis remains insufficiently understood due to the limited genome-level resolution of constituent microbiota. This study addresses this gap by applying genome-resolved metagenomics to connect microbial diversity with agriculture-associated ecological functions in a complex fermented local formulation. By integrating metabolic reconstruction with plant-associated functional traits, this study advances understanding of how microbial consortia contribute to nutrient mobilization, rhizosphere competence, and environmental adaptability. This highlights the contribution of yet-to-be-cultivated but metabolically versatile taxa, which are important to the functions of agricultural ecosystems. By uncovering the roles of key microbes and their cooperative metabolic interactions, this work provides a scientific basis for improving Jeevamrit formulations through informed selection or enrichment of functionally important microbes to enhance nutrient delivery and plant growth performance.

## Introduction

The widespread application of intensive chemical fertilizers has severely compromised global soil health, driving a drastic decline in soil organic carbon, nutrient balance, and microbial diversity (1). To mitigate this critical agricultural challenge, regenerative agriculture practices are gaining significant traction as viable, sustainable methods, with a central focus on biological restoration of soil health (2). In this context, natural farming systems are increasingly adopted to enhance the biogeochemical integrity of soil ecosystems (2, 3). Across the Asian subcontinent, the use of bioformulations such as Jeevamrit and related variants, such as Ghanjeevamrit and Beejamrit (4) are thoroughly used. Jeevamrit is a local fermented liquid bio-inoculant prepared from cow dung, urine, jaggery, pulse flour, and soil, aimed at enriching soil with agri-beneficial microorganisms driving important biochemical processes (5). However, its functional outcomes vary significantly with raw materials and method of preparation, including selection of cattle breed, mixing intensity, environmental exposure, geographic origin of the soil used, etc., thus influencing microbial diversity, nutrient profile, and plant growth attributes (6, 7). Furthermore, variability across preparation cycles, agroclimatic conditions, and lack of mechanization limit the standardization of such formulations (8). Recent metagenomic studies have identified dominance of *Bacillus* and *Pseudomonas* as core community members and reported to impart various growth-promoting attributes (1, 3). The abundance of these taxa is closely linked to physicochemical properties shaped by preparation conditions (6). For instance, continuous aeration enriches *Acinetobacter* (∼40%) and enhances trace metal solubilization, whereas anoxic conditions favor soluble iron and ammoniacal nitrogen accumulation mediated by Bacilli in differentially mixed Jeevamrit samples subjected to a gradient of oxygenation (7). Additionally, enrichment of taxa such as *Bradyrhizobium, Streptomyces, Microvirga, Sphingomonas*, and *Priestia* supports nutrient cycling, improving N, P, and K availability, soil organic carbon, and enzymatic activities, ultimately enhancing crop productivity (2, 4). Application of Jeevamrit modulates crop-specific physiological traits and yields by enhancing nutrient assimilation and reshaping the soil microbiome. In wheat, it has improved biomass and grain yield, doubling the economic cost-benefit ratio (1), while in cabbage, it has boosted top-soil properties and vital enzyme activities, fostering a stable microbial network enriched with beneficial taxa like *Pedomicrobium* and *Solirubrobacter* (5). In turmeric, Jeevamrit and Ghanjeevamrit are reported to enrich beneficial taxa such as *Priestia*, enhancing soil enzyme activities and resulting in over two-fold higher rhizome yield compared to conventional practices (2), thus justifying its role in plant growth promotion.

Though 16S rRNA gene-based metabarcoding has been widely used to characterize microbial diversity and phylogeny, its limited taxonomic resolution and inability to capture the metabolic and functional diversity of the microbiome have led to the use of other genomic methods, such as metagenome-assisted recovery of complete genomes (9). Though cultivation-based studies have highlighted the role of key Jeevamrit members in exerting PGP traits, these members represent only a negligible fraction of the microbiome, given the high and complex microbial load in Jeevamrit. Metagenomic sequencing confirms this vast uncultured majority, revealing Shannon alpha diversity indices ranging from 4.61 to 9.18 and communities averaging 20,684 Operational Taxonomic Units (OTUs) per sample (7, 8). To note, many of the ecologically important taxa remain recalcitrant to cultivation under standard laboratory conditions, although its critical importance to agricultural processes and yield outcomes (10). Overcoming such limitations, metagenome-assembled genomes (MAGs) offer genome-level resolution on microbial identity (uncultivated microbial dark matter) and have substantially expanded our understanding of microbial functionality in the context of agronomic benefit and stress alleviation in crops (11). For instance, MAGs affiliated with *Methylobacterium, Azospirillum*, and *Bradyrhizobium* primarily drive biological nitrogen fixation (12), while specific salinity-associated MAGs affiliated with *Natronomonas* and *Spiribacter* contain genes for plant growth promotion and salinity stress alleviation (13). Genome-level analyses further demonstrate the role of *Amaricoccus, Hydrogenophaga*, and *Brevundimonas* for N-acyl homoserine lactone (AHL)-mediated quorum sensing, central carbon metabolism, and nitrification (14). Functional annotations link MAGs of *Chloroflexota, Armatimonadota*, and *Cyanobacteria* to carbohydrate degradation and biofilm formation (15), and its metabolic connection to *Pseudomonas, Thermoproteota, Actinobacteriota*, and *Desulfobacterota* for enhanced drought resilience (16). Therefore, understanding metagenome-enabled MAGs would help us in identifying critical microbial functions in the context of plant growth and agricultural production.

In the current study, shotgun metagenomic sequencing of two critical Jeevamrit formulations. The selection was based on our previous investigation of Jeevamrit under different mixing conditions that are decisive to the microbial community composition, its association with physicochemical properties (nutrient turnover & dynamics in soil) and plant growth using mainly 16S rRNA gene-based metagenomics (7). Here, we identified a total of 182 bins using binning methods, of which, 16 high-quality MAGs were shortlisted for detailed characterization. Genome-resolved taxonomic and comparative functional analyses were performed to evaluate their metabolic capacity for organic carbon turnover, elemental transformation, habitat colonization, and plant growth promotion traits. In addition, MAGs representing novel lineages were named (sp. nov.) following standard nomenclature rules (SeqCode). The study decoded the multi-layered metabolic synergism of the Jeevamrit’s microbiome and its potential as an effective indigenous microbial inoculant for sustainable agricultural production under low-input methods, i.e., Natural farming.

## Result and Discussion

### Jeevamrit’s metagenomic assembly and recovery of MAGs

The Jeevamrit prepared under constant mixing (CM) promoted aerobic metabolism, enhancing organic matter turnover, oxidative nutrient transformations, and mineral weathering-driven nutrient mobilization. In contrast, no mixing (NM) promoted reductive nutrient transformation, including alternative electron acceptor respiration and fermentative pathways. To resolve the genomic basis of these metabolic regimes, shotgun metagenomics was performed. A total of ∼30.2 million (CM) and ∼37.2 million (NM) filtered reads assembled into ∼891 Mb and ∼1105 Mb of draft metagenomes. Gammaproteobacteria dominated both communities but were more abundant in NM (57.58%) than CM (39.85%), with concurrent enrichment of Bacilli in NM (17.5% vs. 13.14%). CM instead showed higher Bacteroidia (15.88% vs. 11%), Clostridia (13.11% vs. 3.65%), and Betaproteobacteria (11.52% vs. 6.78%). At the genus level, *Acinetobacter* was the most abundant in both CM (31.56%) and NM (41.69%), followed by *Pseudomonas* and *Klebsiella*. CM was further enriched in *Bacteroides* (16.33% vs. 6.74%) and *Comamonas* (11.29% vs. 3.68%), whereas NM had *Trichococcus* (6.54% vs. 2.95%) **(Fig. 1A)**. COG annotations revealed genes assigned to amino acid transport and metabolism (E), energy production and conversion (C), and carbohydrate transport and metabolism (G). CM allocated more genes to information storage and processing than NM (19.66% vs. 18.03%), driven by replication, recombination, and repair (L) and translation (J).

**Fig. 1.**
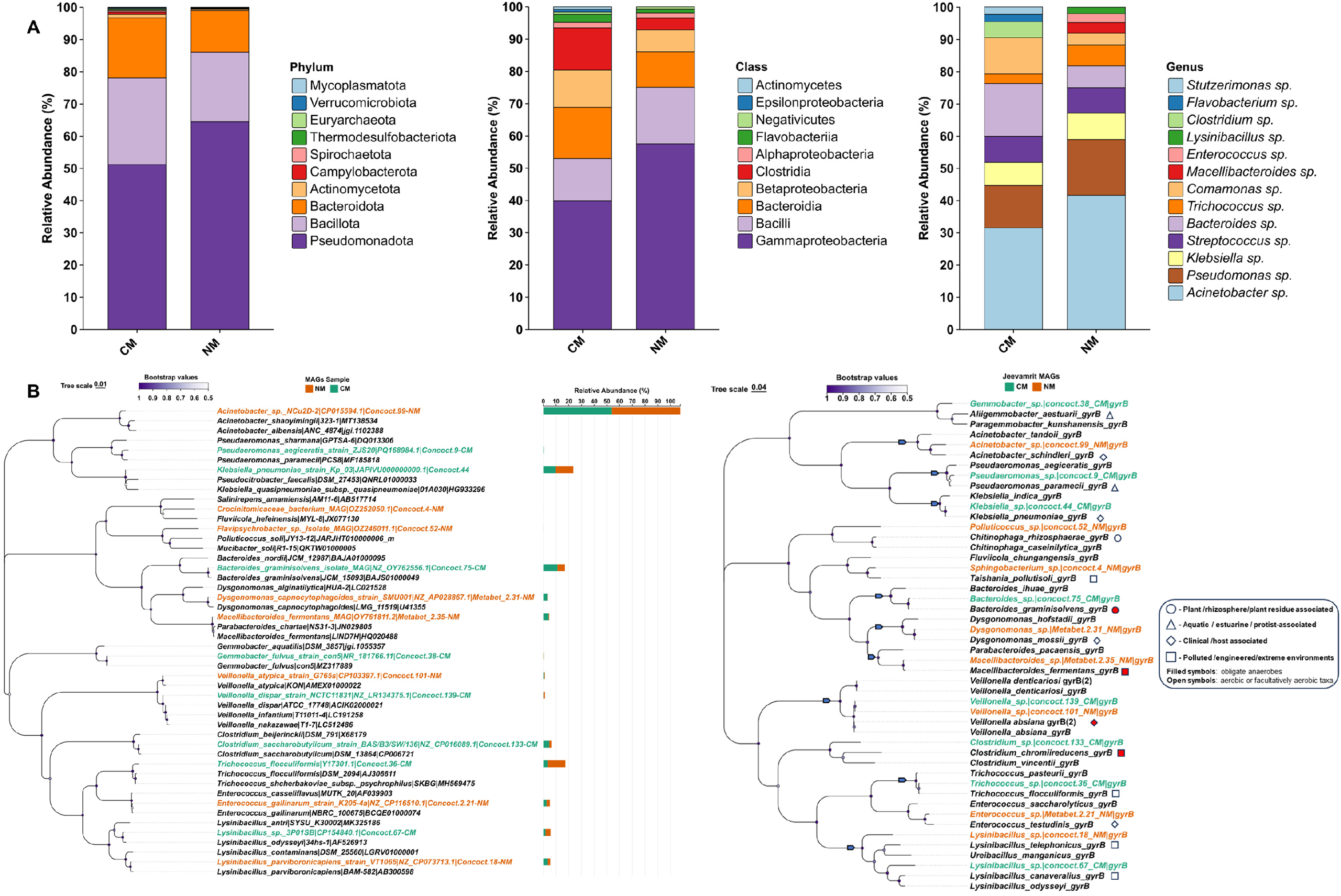
Taxonomic composition and phylogenetic relationships of CM and NM metagenomes. (A) Relative abundance (%) of microbial taxa at phylum, class, and genus levels in conventional management (CM) and natural management (NM) metagenomes based on Kaiju classification. (B) Maximum-likelihood based phylogenetic trees: 16S rRNA gene (left) and *gyr*B (right) with 1,000 bootstrap replicates. MAGs from CM and NM are indicated in orange and green fonts, respectively. Adjacent bars show the relative abundance of each MAG in CM and NM samples.

Metagenomic binning resulted in 182 bins, from which 16 high-quality MAGs (8 CM, 8 NM; >90% completeness, <15% contamination) were recovered, affiliated with Pseudomonadota (n = 5; mean genome size 3.51 Mb), Bacillota (n = 6; 2.72 Mb), and Bacteroidota (n = 5; 3.38 Mb). The genomic attributes were consistent with closest taxonomic relatives. Detailed descriptions of all recovered MAGs, including phylogenetic placement and genome-based relatedness metrics, are provided in the Supplementary file **(Text S1)**. MAG characteristics, including genomic features and relatedness metrics, are summarized in **Table 1**, while detailed comparisons with multiple closest reference taxa and extended genome-based metrics are provided in the Supplementary file **(Table S1)**, and phylogenetic reconstruction based on 16S rRNA and *gyrB* is shown in **Fig. 1B**. GTDB-based phylogenomic analysis corroborated MAG placement across major bacterial lineages **(Fig. S1)**. Considering Jeevamrit’s formulation using farm-based resources (animal origin, such as cow dung), the mapping of antibiotic resistance markers was performed. Consistent presence of glycopeptide resistance genes, mainly partial *van* cluster components (*vanT, vanY, vanW, vanH, vanS*) were noted across most MAGs. Acquired ARGs were scarce, limited to *CMY-4* and *EF-Tu*– associated, and *sul1*/*sul2*, reflecting a lower resistance profile of such taxa. Circular genome maps depicted functional genome architecture, while synteny analysis indicated conserved gene order alongside structural rearrangements **(Fig. S2A, B, S3)**. BGC analysis revealed diverse NRPS, PKS, RiPP, terpene, and β-lactone clusters linked to iron acquisition, antimicrobial activity, quorum sensing, and oxidative stress protection. Several clusters exhibited similarity to known systems, including yersiniabactin-, chrysobactin-, ambactin-, and carotenoid-like pathways, indicating conserved functional potential **(Table S2)**. COG-based gene distributions differed significantly (*p* < 0.05) among categories (Friedman χ^2^ = 247, df = 20, p < 0.001; Kendall’s W = 0.77), with BH-adjusted Wilcoxon tests confirming enrichment of core metabolic and informational functions in these MAGs, including replication and repair (L), amino acid (E) and carbohydrate (G) metabolism, inorganic ion transport (P), energy production (C), and translation (J) **(Fig. S4)**. RAST subsystem analysis also showed significant differentiation (χ^2^ = 107, df = 18, p < 0.001; W = 0.37), highlighting enrichment of central carbon metabolic pathways, particularly anaplerotic and acetogenic pyruvate metabolism, as well as mixed-acid fermentation and associated transporters (ABC family transporters) for uptake of such compounds. Further, detailed functional genomic analyses were performed to infer the phylogenetic demarcation of each MAG, their metabolic inventories, and traits associated with plant growth and niche adaptation/colonization **(Fig. 2)**. PGPT analysis further indicated a diverse distribution of plant growth–promoting traits across MAGs **(Fig. S5)**, as discussed in the following sections.

**Table 1.** Genomic characteristics and taxonomic assignment of metagenome-assembled genomes (MAGs) recovered from CM and NM metagenomes.

| Sample | Bin ID | Genome size (bp) | Completeness (%) | Contamination (%) | Strain heterogeneity | GC (%) | CDS s | Closest relative (TYGS) \| (GTDB) | Phylum | 16S rRNA similarity (%) | OrthoANI (%) | dDDH (%) | ΔG+C (%) | Predicted Functional Role in Agroecosystem |
| --- | --- | --- | --- | --- | --- | --- | --- | --- | --- | --- | --- | --- | --- | --- |
| CM | Concoct.9 | 3307077 | 96.08 | 5.89 | 14.29 | 61.89 | 3550 | *Pseud aeromonas paramecii* JCM 32226 | Pseudomonadota | 98.57 | 90.29 | 40.5 | 0.15 | PGP + NU + BC |
|  | Concoct.38 | 2677557 | 91.06 | 4.39 | 66.67 | 58.71 | 3313 | *Gemmobacter aquatilis* D SM 3857 (GTDB: *Pseudotabrizicola* sp.) |  | 97.88 | 72.76 | 19.3 | 6.39 | PGP + NU + BC |
|  | Concoct.44 | 6151882 | 98.94 | 5.03 | 65.43 | 56.61 | 6417 | *Klebsiella pneumoniae* subsp. *pneumoniae* DSM 30104 |  | 99.79 | 99.00 | 91.6 | 0.14 | PGP + NU + BC |
|  | Concoct.36 | 2693805 | 94.06 | 2.93 | 14.29 | 46.7 | 2894 | *Trichococcus flocculiformis* DSM 2094 | Bacillota | 100.00 | 97.34 | 77.6 | 1.03 | PGP + NU |
|  | Concoct.67 | 3514077 | 93.86 | 14.7 | 16.98 | 37.22 | 3913 | *Lysinibacillus odyseyi* NBRC 100172 (GTDB: *Solibacillus* sp.) |  | 99.47 | 71.95 | 20.2 | 3.74 | PGP + NU + BC |
|  | Concoct.133 | 3086410 | 99.19 | 1.69 | 0 | 28.94 | 3404 | *Clostridium cagae* Marseille-P4344 |  | 96.69 | 71.42 | 19.8 | 1.69 | PGP + NU |
|  | Concoct.139 | 1880027 | 98.86 | 0.1 | 0 | 39.99 | 1807 | *Veillonella absiana* YH-vei2232 |  | 99.80 | 99.37 | 94.4 | 0.12 | PGP + NU |
|  | Concoct.75 | 3673511 | 99.25 | 1.28 | 14.29 | 41.51 | 3273 | *Bacteroides graminisolvens* DSM 19988 | Bacteroidota | 99.80 | 97.73 | 84.8 | 0.02 | PGP + NU |
| NM | Concoct.99 | 2710978 | 95.34 | 2.3 | 39.13 | 40.97 | 2620 | *Acinetobacter townieri* DSM 14962 | Pseudomonadota | 93.96 | 77.1 | 22.1 | 0.38 | PGP + NU + BC |
|  | Concoct.18 | 3018196 | 97.35 | 1.21 | 0 | 36.73 | 2953 | *Lysinibacillus sphaericus* KCTC 3346 | Bacillota | 98.62 | 73.33 | 21.6 | 0.39 | PGP + NU + BC |
|  | Concoct.101 | 1932039 | 100 | 0 | 7.14 | 40.01 | 1862 | *Veillonella absiana* YH-vei2232 |  | 99.41 | 99.39 | 94.5 | 0.1 | PGP + NU |
|  | Metabet2.21 | 2904910 | 98.44 | 1.12 | 50 | 40.84 | 2875 | *Enterococcus gallinarum* NBRC 100675 |  | 99.81 | 97.69 | 80.2 | 1.09 | PGP + NU |
|  | Concoct.4 | 3349813 | 100 | 13.46 | 10 | 36.02 | 3194 | *Sphingobacterium cavernae* 5.0403-2 (GTDB: *Taishania* sp.) | Bacteroidota | 84.10 | 75.68 | 70.1 | 0.22 | PGP + NU + BC |
|  | Concoct.52 | 3264722 | 92.69 | 5.79 | 100 | 34.33 | 3529 | *Polluticoccus soli* JY13-12 (GTDB: *Edaphocola* sp.) |  | 92.56 | 66.06 | 19.5 | 11.84 | PGP + NU + BC |
|  | Metabet2.31 | 3566855 | 95.36 | 3.42 | 66.67 | 37.48 | 3392 | *Dysgonomonas mossii* DSM 22836 |  | 92.14 | 96.13 | 69.3 | 0.01 | PGP + NU + BC |
|  | Metabet2.35 | 3054732 | 93.65 | 2 | 40 | 42.15 | 2800 | *Macellibacteroides fermentans* DSM 24967 |  | 99.80 | 98.36 | 83.5 | 0.27 | PGP + NU + BC |

**Fig. 2.**
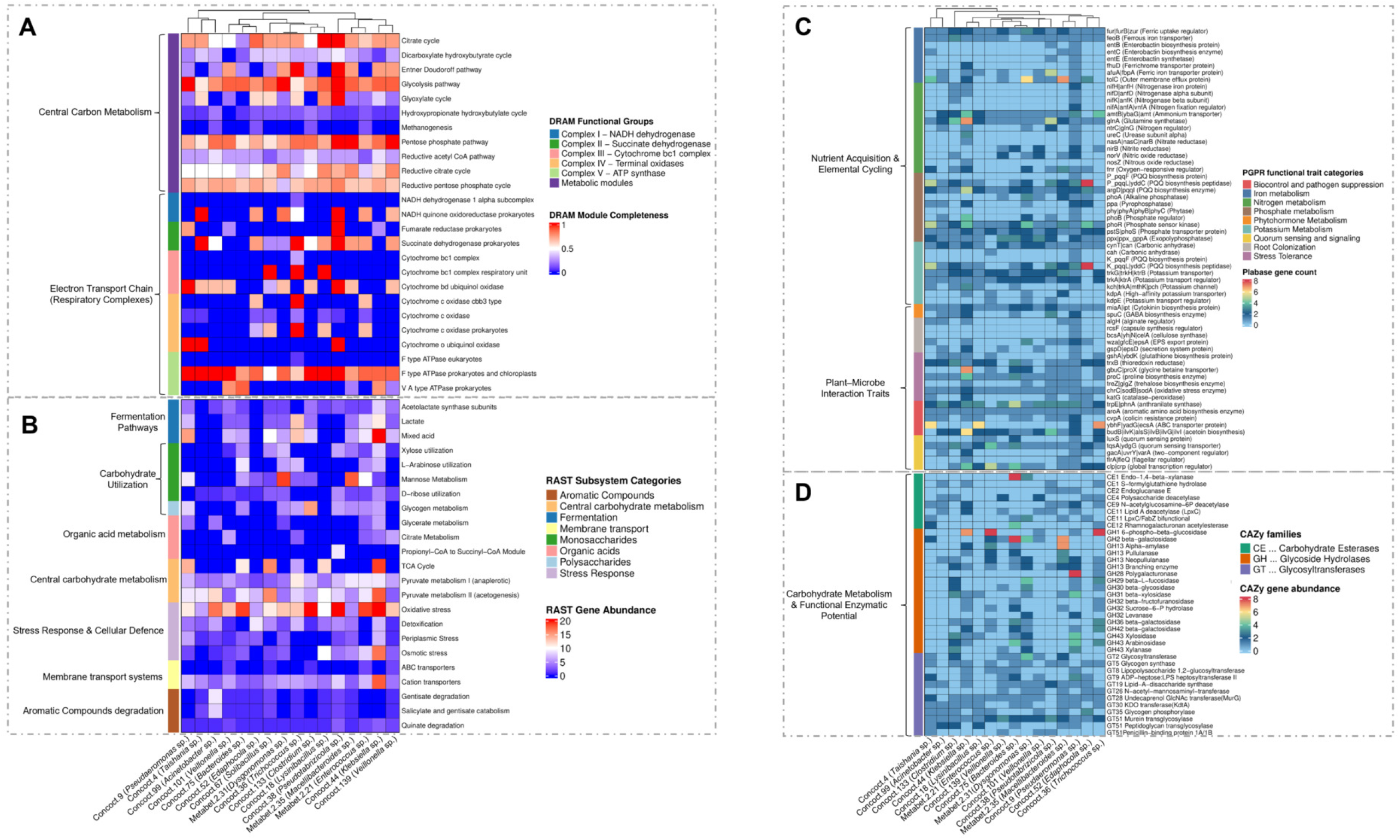
Genome-resolved functional potential (heatmap) of 16 MAGs recovered from CM and NM metagenomes. (A) DRAM-annotated functional groups showing metabolic module completeness across MAGs, including central carbon metabolism, fermentation pathways, and respiratory complexes (Complex I–V). The color scale represents module completeness. (B) RAST subsystem categories depict gene abundance across major functional groups, including carbohydrate metabolism, amino acid metabolism, stress response, membrane transport, and aromatic compound degradation. (C) Plant growth–promoting and nutrient cycling traits based on PLaBAse annotation, including Fe, N-P-K metabolism, phytohormone production, root colonization, stress tolerance, biocontrol, and quorum sensing. The color scale represents gene counts within each MAG. (D) Distribution of carbohydrate-active enzyme (CAZy) families across MAGs, including carbohydrate esterases (CEs), glycoside hydrolases (GHs), and glycosyltransferases (GTs). Color intensity reflects gene abundance

#### (i) Pseudomonadota

A total of five MAGs were recovered within Pseudomonadota and were affiliated with *Pseudaeromonas, Gemmobacter, Klebsiella*, and *Acinetobacter. Acinetobacter* MAG was detected exclusively in NM, whereas the remaining MAGs were also associated with CM.

##### (a) Concoct.9 (*Pseudaeromonas* sp.)

Phylogenetically closest to *P. paramecii* JCM 32226 (16S rRNA 98.57%; MLSA ∼98.0%; GTDB classifier), with genome-based indices (ANI 90.29%, dDDH 40.5%) supporting its designation as a novel species, for which Candidatus *Pseudaeromonas jeevamritensis* sp. nov. is proposed (jee.va.mri.ten’sis; N.L. adj., explaining its abundance in Jeevamrit). Carbon metabolism genes encode monosaccharide utilization (especially mannose), glycogen hydrolysis, mixed-acid fermentation (acetate and D-lactate), organic acid utilization, and pyruvate anaplerotic routes (for acetyl-CoA and formate). Multiple glycosyltransferases (GT8, GT9, GT19, GT28, GT30, GT51), carbohydrate esterases (CE2, CE9), and glycoside hydrolase (GH1) families validate carbohydrate processing capacity, consistent with *Pseudaeromonas* versatility in oxidizing sugars, hydrolyzing starch, and conducting facultative anaerobic fermentation to produce acids without gas (17). A complete Fe-acquisition system (*fur, feoB, entBEC, fhuD, afuA, tolC*) for iron regulation/uptake, *nifHDK* with regulator *nifA* for N-fixation, and redox assimilation genes (*amtB, glnA, nasA, nirB, norV*) were detected. Phytohormone genes (*amiE, miaA/ipt, spuC*), P-metabolism (*pqqFL, phoABR, pstS, ppx*), and K-transport (*trk, kdp*) support nutrient acquisition and ionic homeostasis. Its closest relative, *P. paramecii* JCM 32226, similarly harbors these N-fixation, P-response, and auxin biosynthesis systems, confirming plant-beneficial properties (18). Genes related to biofilm formation and colonization (*algH, rcsF, bcsA, wza, gspD*), oxidative and osmotic stress tolerance (*sodB, katG, gshA, trxB, gbuC, proC, treZ*), biocontrol functions (*trpE, aroA, cvpA, ybhF, budB*), quorum-sensing and global regulation (*luxS, gacA, fleQ, crp*), and transporter (*tqsA*) were noted. Based on these genomic traits, this taxon likely plays a key role in Jeevamrit’s microbiome through heterotrophic carbon turnover linked to N-transformation, with environmental survival and cell-cell communication capabilities.

##### (b) Concoct.38 (*Pseudotabrizicola* sp.)

Phylogenetically closest to *Gemmobacter aquatilis* (16S rRNA 97.88%; MLSA ∼91.6%), while GTDB assigns it to *Pseudotabrizicola*. Genome-based indices (ANI 72.76%, dDDH 19.30%) support its designation as a novel species, for which Candidatus *Pseudotabrizicola heterotrophica* sp. nov. is proposed (he.te.ro.tro’phi.ca; N.L. fem. adj. heterotrophica, referring to a heterotrophic mode of nutrition). Genome mining revealed oxidative pathways for organic C-metabolism, including glycolysis, xylose and glycogen metabolism, acetolactate synthesis, and propionyl-CoA-to-succinyl-CoA conversion coupled to the aerobic ETC for energy generation. CAZy annotation showed that carbohydrate esterases (CE2, CE9, CE11) and glycosyltransferases (GT19, GT26, GT51) are the major enzymes involved in envelope biogenesis rather than polysaccharide degradation. The MAG contained N-assimilation genes with regulators and transporters (*amtB, glnA, ntrC*). Genes for phytohormone synthesis (*amiE, miaA/ipt, spuC*), root colonization (*algH, wza*), stress tolerance (*gshA, trxB, sodB, katG*), and biocontrol (*trpE, aroA, cvpA, budB*) were identified. Comparative genomics reveals diverse metabolic traits in close relatives: Gemmobacter taxa exhibit methylated amine utilization linked to C–N cycling (*G. aquatilis* DSM 3857) (19). *Pseudotabrizicola* members harbor aerobic anoxygenic photosynthesis (*P. sediminis, P. algicola*) and alkaline adaptation (*P. alkalilacus*), and occur in organic-rich aquatic habitats, including saline sediments, alkaline waters, and microalgal consortia (20,21). The recovery of Concoct.38 from Jeevamrit expands niche reporting for this lineage and suggests broader functional potential through phytohormone synthesis, root colonization, and biocontrol capabilities.

##### (c)Concoct.44 (*Klebsiella* sp.)

Demonstrates the closest genomic relatedness to *Klebsiella pneumoniae* subsp. *pneumoniae* DSM 30104. Metabolic reconstruction indicated broad carbon utilization (xylose, L-arabinose, mannose, glycogen, citrate) and mixed-acid fermentation capacity, including EMP and ED pathways, a complete glyoxylate shunt, and multiple anaplerotic modules. Enrichment of glycoside hydrolases (GH1, 13, 31, 32, 36, 42, 43) and glycosyltransferases (GT2, 5, 8, 19, 26, 30) validates this metabolic breadth. The MAG encodes iron uptake and regulation (*fur, feoB, fhuD, afuA, tolC*), N-fixation (*nifHD*), assimilation (*amtB, glnA, nasA, fnr*), and urease (*ureC*) for enhanced N turnover. Phytohormone production, root colonization, biofilm formation, and stress resilience supported by osmoprotectant transporters (*gbuC/proX*) and redox enzymes (*gshA, trxB, proC, sodB, katG*) were identified. The highest PGPT count (6,769) among all MAGs indicated superior plant growth-promoting capacity **(Fig. S5)**. The occurrence of this MAG in Jeevamrit aligns with *Klebsiella* prevalence in soil, water, and plant-associated habitats, supported by dynamic nutrient biotransformation. Members exhibit well-documented PGP traits, including N-fixation, phytohormone production, Fe-acquisition, and P-solubilization (22). *K. pneumoniae* 342 fixes N_2_ in wheat, contributing approximately 37-49% of plant nitrogen from atmospheric sources under N-deficient conditions (23). Similar traits are reported in *Klebsiella* from Panchagavya (fermented cow-based formulations), supporting roles in C-N cycling and rhizosphere competence (24).

##### (d) Concoct.99 (*Acinetobacter* sp.)

Most closely related to *Acinetobacter towneri* DSM 14962 based on 16S rRNA gene, with GTDB supporting *Acinetobacter* affiliation. Genome-based indices (ANI 77%, dDDH 22%) support its recognition as a novel species, for which Candidatus *Acinetobacter phytophilus* sp. nov. is proposed (phy.to’phi.lus; N.L. masc. adj., plant-associated or plant-loving). Notably, *Acinetobacter* spp. were relatively abundant in both NM and CM samples **(Fig. 1B)**, suggesting enrichment independent of mixing regimes and, based on abundance, a potentially important role in Jeevamrit functional outcomes. The presence of Concoct.99 aligns with the genus’s metabolic versatility and wide distribution in soil and aquatic environments (25). Salicylate and gentisate degradation pathways, pyruvate-driven acetogenesis, and a complete glyoxylate shunt support efficient carbon turnover in complex niches (26). Abundant glycosyltransferases with limited glycoside hydrolases suggest emphasis on cell envelope formation rather than extensive polysaccharide degradation (27). Genes associated with phytohormone production (*amiE, miaA/ipt*) and colonization (*algH, gspD*) indicate PGP potential, consistent with documented phytohormone-mediated growth promotion and rhizosphere colonization in *Acinetobacter* spp. (28, 29). Close relatives demonstrate relevant agricultural functions: *A. towneri* accumulates up to 24.57% (w/w as P) intracellular polyphosphate under sulfur limitation, enabling P-sequestration (30); *A. schindleri* SR-5-1 enhances plant growth, antioxidant activity, nutrient acquisition, and reduces Cd accumulation under wastewater/Cd stress (31); and strain H4-3-C1 degrades *Fusarium* toxins (e.g., zearalenone), supporting plant-protective endophytism (32). Although N-fixation is absent, N-assimilation regulators (*amtB*), Fe-acquisition systems (*feoB*), and stress tolerance genes (*gshA, sodB*) highlight this MAG’s potential role in nutrient acquisition and environmental adaptation.

#### (ii) Bacillota

Six Bacillota-affiliated MAGs representing *Trichococcus, Lysinibacillus, Clostridium, Veillonella*, and *Enterococcus* were recovered, with clear partitioning between CM (*Trichococcus, Lysinibacillus, Clostridium*) and NM (*Lysinibacillus, Veillonella, Enterococcus*).

##### (a) Concoct.36 (*Trichococcus* sp.)

Closest to *Trichococcus flocculiformis* DSM 2094 based on 16S rRNA and *gyrB* similarity, supported by genome phylogenomics. Metabolic reconstruction indicated fermentative C-metabolism with pyruvate-to-lactate conversion and mixed-acid by-products (acetate, formate, ethanol), supported by glyoxylate and reductive acetyl-CoA modules. The presence of glycoside hydrolases (GH13, GH23, GH32), glycosyltransferases (GT2, GT4), and carbohydrate esterases (CE4) might indicate its polysaccharide turnover and biosynthesis. Closely related to *T. flocculiformis* DSM 2094, this taxon carries genomic determinants for biofilm formation, capsule production, and adhesion for environmental residence (33). The presence of cold shock proteins and osmoprotectant transporters indicates tolerance to low temperature and salinity stress (34). The presence of Concoct.36 in Jeevamrit aligns with the genus’s role as a metabolically versatile member of anaerobic and facultative environments.

##### (b) Concoct.67 (*Solibacillus* sp.)

Phylogenetically closest to *Lysinibacillus odysseyi* (16S rRNA similarity 99.47%); however, GTDB assigns it to *Solibacillus*. Genome-based indices (ANI 71%, dDDH 19%) support its recognition as a novel species, for which Candidatus *Solibacillus xylolyticus* sp. nov. is proposed (xy.lo.ly’ti.cus; N.L. masc. adj., referring to xylose-degrading capacity). Metabolically, Concoct.67 encodes xylose utilization and a cyt bc_1_ complex, indicating active respiratory metabolism. Its CAZy profile shows limited glycoside hydrolases (notably GH2) but enrichment of glycosyltransferases (GT2, GT5, GT8, GT9, GT35, GT51), suggesting preference for cell wall-associated glycan biosynthesis rather than extensive polysaccharide degradation. The MAG encodes Fe-homeostasis, N-assimilation, P-metabolism, and K-transport systems for nutrient acquisition. Genes related to phytohormone modulation (*amiE, miaA/ipt, spuC*), biocontrol and quorum sensing (*trpE, aroA, cvpA, budB, luxS, crp, gacA, tqsA*) indicate antimicrobial potential and rhizosphere fitness. Comparative genomics reveals diverse functional traits in close relatives: *L. odysseyi* Gr102 produces antimicrobial metabolites active against pathogens, while *L. xylanilyticus* exhibits xylan-degrading capacity (35). *Solibacillus* taxa demonstrate extreme environmental resilience, including survival in the upper atmosphere and on the International Space Station (36), Fe-chelating siderophore production in soil isolates, and the production of complex extracellular compounds, such as amyloidogenic biosurfactants and bioflocculants (37). The recovery of Concoct.67 from Jeevamrit links the genus’s robust stress tolerance, xylose utilization, and capacity to produce protective extracellular compounds, enabling adaptation to alkaline, nutrient-rich, and highly competitive environments.

##### (c) Concoct.18 (*Lysinibacillus* sp.)

Concoct.18 is affiliated with *Lysinibacillus sphaericus* (16S rRNA similarity 98.62%), and genome-based indices (ANI 73%, dDDH 21%) support its designation as a novel species, for which the name Candidatus *Lysinibacillus redoxflexus* sp. nov. is proposed (re.dox.flex’us; N.L. masc. adj., referring to metabolic flexibility under varying redox conditions). Concoct.18 encodes multiple reductive metabolic modules, including the reductive tricarboxylic acid (rTCA) cycle, reductive pentose phosphate pathway (rPPP), and dicarboxylate-hydroxybutyrate cycle, indicating fluctuating redox conditions. The CAZy profile includes glycoside hydrolases (GH31, GH43) along with enriched glycosyltransferases (GT26, GT28, GT35, GT51), suggesting a balance between carbohydrate processing and structural glycan biosynthesis. It also encodes systems for Fe homeostasis (*fur, feoB*), with the presence of *afuA* enhancing ferric iron uptake. Nutrient cycling capabilities include N-assimilation (*amtB, glnA, nirB*), P-metabolism (*pqqL/I, phoB, phoR, pstS, ppx*), and K-transport (*trk, kdp*). Phytohormone-related genes (*amiE, miaA/ipt*) indicate plant interaction potential, while conserved stress-response genes (*gshA, sodB*) contribute to oxidative stress tolerance. Biocontrol and signaling genes (*trpE, aroA, cvpA, budB, luxS, crp*) highlight its ecological competence in the rhizosphere. Supporting evidence from related strains shows that *L. sphaericus* ZA9 produces high levels of IAA, siderophores, HCN, and hydrolytic enzymes, and enhances crop yields, while *L. sphaericus* KCTC 3346 harbors NRPS/PKS clusters for secondary metabolite biosynthesis (38). Although these traits are consistent with known *Lysinibacillus* functions, the presence of distinct reductive pathways (rTCA, rPPP) and the specialized CAZy profile observed in Concoct.18 could signify a spore-forming, redox-active metal biotransformation (39) and agriculturally important secondary metabolites production-based genomic traits.

##### (d) Concoct.133 (*Clostridium* sp.)

Closest to *Clostridium gasigenes* DSM 12272, with both 16S rRNA and GTDB placing it within *Clostridium*. Genome-based indices (ANI 71%, dDDH 19%) support its designation as a novel species, for which Candidatus *Clostridium fermentans* sp. nov. is proposed (fer.men’tans; L. pres. part., fermenting, referring to strong organic acid–producing metabolism). Metabolic reconstruction indicates fermentative metabolism with lactate and mixed-acid production, supported by reductive acetyl-CoA, rTCA, rPPP, and dicarboxylate–hydroxybutyrate modules, reflecting anaerobic flexibility. CAZy profiles comprise carbohydrate esterases (CE4), glycoside hydrolases (GH13, 29, 30, 31, 36, 42, 43), and glycosyltransferases (GT2, 26, 28, 30, 35, 51). Membrane transport is dominated by ABC and cation transporters. It encodes N-fixation (*nifH, nifD*) and assimilation regulators (*amtB, glnA, fnr*), supporting N-transformation. Fe-homeostasis (*fur, feoB*), P-metabolism (*pqqL, phoB, phoR, pstS, ppx*), K-transport (*trk, kdp*), and phytohormone genes (*amiE, miaA/ipt, spuC*) suggest nutrient acquisition and growth-modulation abilities. The isolation of Concoct.133 from Jeevamrit aligns with *Clostridium* prevalence in manure, soil, and nutrient-rich anaerobic environments, where diverse glycoside hydrolases enable intensive turnover of complex carbohydrates, including starch and crystalline cellulose. Fermentation products such as butyrate and lactate maintain biological homeostasis (40). Close relatives demonstrate relevant metabolic capacities: *C. chromiireducens* conducts mixed-acid fermentation (acetate, butyrate, formate, lactate) coupled to Cr (VI) reduction under anaerobic conditions (41), while *C. saccharobutylicum* converts poplar hydrolysates to ABE solvents, yielding up to 6.98 g/L butanol and 9.64 g/L ABE (42). These ABE-solvent and reductive pathway capabilities indicate potential for anaerobic carbon flux and nutrient turnover, enhancing nutrient availability and plant health in fermented systems (43).

##### (e) Concoct.139 and Concoct.101 (*Veillonella* spp.)

Both show the strongest affinity to *Veillonella absiana* YH-vei2232 based on 16S rRNA (99.80% and 99.41%) and MLSA (∼98%). Both MAGs exhibit anaerobic fermentation with prominent reductive modules (reductive acetyl-CoA, rTCA, rPPP; with dicarboxylate-hydroxybutyrate in Concoct.101). Energy conservation features fumarate reductase (Concoct.139) and succinate dehydrogenase (Concoct.101). CAZy repertoires are limited in degradative enzymes (GH1, 2, 13, 30; CE4, 11, 12) but enriched in glycosyltransferases (GT2, 5, 26, 28, 30, 35, 51), emphasizing glycan modification and cell envelope biosynthesis, a specialized metabolic adaptation enhancing fermentation flexibility not yet detailed in the literature. Both MAGs encode Fe-acquisition (*fur, feoB, afuA, tolC*), N-assimilation/reduction (*amtB, glnA, nirB*; *nasA* in Concoct.101; *fnr* in Concoct.139), P-sensing/transport (*phoB, phoR, pstS, ppx*; *pqqL* in Concoct.139), and K-transport (*trk, kdp*). Phytohormone genes (*amiE, miaA/ipt, spuC*) and colonization/secretion determinants (*gspD*; *wza* in Concoct.139) indicate plant interaction. Stress and regulatory modules (*gshA, sodB, trxB, katG*) with biocontrol genes (*trpE, aroA, cvpA, budB*) and QS regulators (*luxS, gacA, crp*) support oxidative resilience and competitive fitness. The presence of these MAGs in Jeevamrit reflects the occurrence of *Veillonella* in the animal gut microbiota (44) and its adaptation to the anaerobic, nutrient-rich conditions created by cattle dung and urine. The genus is characterized by a “Gram-negative Firmicutes” profile and functions as a secondary fermenter, producing acetate and propionate from lactate (45). N-assimilation and partial nitrate/nitrite reduction pathways support a role in nitrogen turnover under reduced conditions and in maintaining metabolic flux within complex anaerobic communities.

##### (f) Metabet 2.21 (*Enterococcus* sp.)

It exhibits the closest relationship to *Enterococcus gallinarum* based on 16S rRNA (99.81%) and MLSA (∼95–96%). Functional annotation indicates fermentative heterotrophy with mixed-acid and lactate production as the major central carbon pathway supported by reductive PPP, rTCA, and reductive acetyl-CoA modules. Energy conservation involves fumarate reductase, cytochrome bd oxidase, and F/V-type ATP synthases. The presence of Metabet 2.21 in Jeevamrit reflects the genus’s ability to persist in environmental reservoirs, such as soil, manure, and fermented ecosystems (46). Its presence in Jeevamrit aligns with the genus’s role in early-stage fermentation and tolerance to anaerobic, nutrient-rich manure environments (47). This isolate is taxonomically affiliated with the *E.gallinarum* lineage, an efficient fermentative heterotrophy (48). The P-solubilization, and plant growth–promoting traits were identified in this MAG correspond with reports describing *Enterococcus* species as halotolerant rhizobacteria that enhance nutrient mobilization and crop yield (49). Additionally, the antifungal potential and stress-related genes align with *Enterococcus’s* role in pathogen suppression and oxidative stress tolerance in the rhizosphere (50). While these PGP traits are known, the presence of GH32 and GH36 with reductive modules suggests specialized carbohydrate utilization that has not been well explored in *Enterococcus*. Collectively, these features highlight the isolate’s potential to promote plant growth through organic acid-mediated nutrient turnover and improved competitive fitness

#### (iii) Bacteroidota

Five Bacteroidota MAGs were recovered, affiliating with *Bacteroides* from CM and *Sphingobacterium, Polluticoccus, Dysgonomonas*, and *Macellibacteroides* from NM.

##### (a) Concoct.75 (*Bacteroides* sp.)

Displays closest affinity to *Bacteroides graminisolvens* DSM 19988 based on genome phylogenomic and *gyrB*. Functional reconstruction indicates a mixed-acid fermentative metabolism with auxiliary reductive modules (rTCA, dicarboxylate-hydroxybutyrate). The genome supports utilization of xylose, L-arabinose, mannose, and aromatic compounds (quinate, salicylate, gentisate), with cytochrome bd oxidase supporting respiration-linked flexibility. CAZy annotation revealed glycoside hydrolase families (GH1, 2, 13, 32, 36, 42, 43) and glycosyltransferase families (GT2, 5, 8, 9, 19, 26, 28, 30, 35, 51), supporting simple sugar metabolism and glycoconjugate assembly. Genomic analysis further indicates traits associated with nutrient acquisition and rhizosphere adaptation, including genes involved in ammonium assimilation (*amtB, glnA*), P-sensing (*phoB*), and high-affinity k-uptake (*kdpA*). These features align with the broader role of Bacteroidetes in nutrient cycling and plant-associated ecosystems (51,52). Phytohormone genes (*amiE, miaA/ipt, spuC*), stress determinants (*gshA, trxB, proC, treZ, sodB, katG*), and biocontrol and regulatory modules (*trpE, aroA, cvpA, budB; luxS, gacA, flrA, crp*) indicate plant interaction, resilience, and competitive fitness. The reference taxon of the MAG, *B.graminisolvens*, is a strictly anaerobic, xylanolytic bacterium isolated from plant residue in a methanogenic reactor that ferments xylan and related saccharides to acetate, propionate, and succinate (53), underscoring its role in hemicellulose degradation and carbon turnover under anaerobic conditions.

##### (b) Concoct.4 (*Taishania* sp.)

Closely related to *Sphingobacterium cavernae* 5.0403-2 based on genome analysis, but assigned to *Taishania* by GTDB. Genome-based indices (ANI 75%, dDDH 70.1%, borderline) support its recognition as a novel species, Candidatus *Taishania aromaticivorans* sp. nov. (a.ro.ma.ti.ci.vo’rans; N.L. part. adj., referring to an organism capable of degrading aromatic compounds). Functional reconstruction indicates aerobic energy metabolism supported by respiratory complexes (I, II, IV) and F-type ATP synthase, as well as aromatic compound degradation (quinate, salicylate, gentisate) and diverse ABC/cation transporters. CAZy profiles include carbohydrate esterases (CE9, CE11) and glycoside hydrolase (GH2), alongside glycosyltransferases (GT2, 8, 19, 28, 30, 35, 51), emphasizing glycoconjugate and envelope biosynthesis over extensive polysaccharide degradation. Genes for N-assimilation and P-solubilization support nutrient acquisition, while stress-response genes indicate oxidative and osmotic tolerance. Biocontrol traits and iron regulation support pathogen suppression, including against *Fusarium* (54). Comparative genomics shows that reference taxa (*S. cavernae* and *T. pollutisoli*) exhibit phosphatases that support organic P-mineralization (55, 56) and rhizosphere-associated, plant growth-promoting functions (56). Members of the taxonomic family Crocinitomicaceae, including *Taishania*, demonstrate ecological resilience across diverse habitats ranging from forest soils, wetlands, and wastewater to marine ecosystems and long-term contaminated soils (56). The recovery of Concoct.4 from Jeevamrit aligns with this broad environmental distribution and suggests potential roles in soil adaptation and restoration.

##### (c) Concoct.52 (*Edaphocola* sp.)

Based on 16S rRNA analysis (92.56%), it is most closely related to *Polluticoccus soli* JY13-12, whereas GTDB places it within *Edaphocola*; genome-based indices (ANI 66.06%, dDDH 19.50%) indicate that it represents a novel species, for which Candidatus *Edaphocola rhizoadaptatus* sp. nov. is proposed (rhi.zo.a.dap.ta’tus; N.L. masc. adj., referring to an organism adapted to the plant rhizosphere environment). Functional reconstruction indicates respiration-supported heterotrophy with reductive modules (glyoxylate, rPPP, rTCA, dicarboxylate-hydroxybutyrate) and a cbb3-type oxidase, suggesting low-oxygen adaptation. A restricted CAZy profile (CE4, CE11; GT families) lacking GHs indicates minimal carbohydrate depolymerization. Genes for P-acquisition, K-transport, and *miaA*/*ipt* support nutrient uptake and plant interaction, while stress, biocontrol, and regulatory genes indicate resilience and ecological adaptability. The detection of Concoct.52 in Jeevamrit highlights its role in supporting nutrient turnover and plant growth under fluctuating redox conditions. The genus *Edaphocola*, recently established from wetland soil, represents a distinct lineage within the family Chitinophagaceae based on phylogenetic, phenotypic, and chemotaxonomic characteristics (57). Most notably, the genus demonstrates a profound capacity for xenobiotic bioremediation; for instance, *E. flava* was isolated directly from herbicide-contaminated soils and utilizes novel esterases (such as *LanE*) to actively degrade complex, chiral herbicides, such as lactofen (58).

##### (d) Metabet 2.31 (*Dysgonomonas* sp.)

Displays closest phylogenomic affiliation with *Dysgonomonas mossii* DSM 22836, supported by *gyrB* similarity. Functional reconstruction indicates a carbohydrate-driven metabolism with mixed-acid/lactate fermentation and auxiliary reductive modules (rTCA, glyoxylate, dicarboxylate-hydroxybutyrate), supported by the genes for utilization of xylose, L-arabinose, mannose, ribose, and glycogen, with cytochrome bd oxidase-linked respiratory flexibility. CAZy profiles include carbohydrate esterase CE1, glycoside hydrolases (GH1, 2, 13, 29, 30, 31, 32, 36, 42, 43), and glycosyltransferases (GT2, 8, 28, 30, 35, 51), supporting hemicellulose and oligosaccharide turnover alongside glycoconjugate biosynthesis. The genome encodes Fe-regulation (*fur, feoB, tolC*), NH_4_^+^ assimilation/regulation (*amtB, glnA, ntrC, fnr*), and a comprehensive P-acquisition system (*pqqL, argD/pqqI, phoA, ppa, phoB/R, pstS, ppx, cynT*), with K-transporters (*trkG/H/ktrB, trkA, kch, kdpA*) supporting ionic balance. *D. mossii* encodes xylan-targeting PULs with CE1/CE6 esterases and GHs that deconstruct decorated hemicellulose, releasing fermentable sugars that are subsequently converted into SCFAs (59). *D. capnocytophagoides* exhibits phosphatase and glycosidase activities, suggesting a role in carbohydrate metabolism (60). Metabet 2.31 likely contributes to hemicellulose degradation and fermentative carbon flux, as Jeevamrit. *D. mossii* exhibits facultative anaerobic and carbohydrate-utilizing capabilities, indicating potential involvement in fermentative carbon processing. Collectively, these genes highlight the isolate’s potential to drive carbon flux and enhance nutrient availability in organically amended soils.

##### (e) Metabet 2.35 (Macellibacteroides sp.)

Closest to *Macellibacteroides fermentans* DSM 24967 based on 16S rRNA (99.80%) and MLSA (∼99.6%), supported by phylogenomics. Genomic reconstruction indicated a saccharolytic, heterotrophic mechanism utilizing xylose, mannose, ribose, and glycogen, with fermentation of mixed acids, lactate, and short-chain fatty acids (acetate and butyrate). Carbohydrate esterases (CE2, CE11) and glycoside hydrolases (GH2, 13, 28, 29, 31, 32, 36, 43), including GH43 and GH13, enable the deconstruction of plant-derived polysaccharides, driving carbon flux and supporting microbial cross-feeding within the formulation. The presence of Metabet.2.35 in Jeevamrit aligns with the genus’s established role as an obligately anaerobic fermenter commonly associated with wastewater treatment systems, including anaerobic filters and sludge-based engineered environments (61). Despite the genus’s obligately anaerobic characterization, this MAG uniquely encodes cytochrome bd oxidase, reductive modules (rTCA), and NH_4_^+^ assimilation (*amtB, glnA*), suggesting specialized metabolic flexibility and N-regulation. The genome encodes Fe-homeostasis (*fur, feoB, entC*; multiple *tolC*), NH_4_^+^ assimilation/regulation (*amtB, glnA, fnr*), and P and K acquisition systems (*pqqL, argD/pqqI, phoA, phoB/R, pstS, ppx, cynT, trkG/H/ktrB, trkA, kch, kdpA*), indicating nutrient mobilization capacity. Combined with biocontrol modules, this underscores the potential of this lineage to enhance rhizosphere resilience and stimulate root development.

It is noteworthy that, based on the 16 MAGs’ genetic compositions, we hypothesize that the Jeevamrit microbiome is highly structured along a gene-centric metabolic and eco-physiological spectrum. Based on composite Z-scores, genomes were classified into three tiers: high (Z = 41.1–124.7), moderate (Z = 9.9– 23.3), and low (Z = −8.4 to −58.8), representing a gradient of compositional and metabolic versatility **(Table S3)** relevant to soil nutrient conditioning and plant growth promotion (PGP). The structured metabolic organization of MAGs reflects a cooperative framework underlying its functional and PGP activity. The upper layer with a high Z-score functions as a primary degrader and nutrient transformer, driving OC turnover and converting substrates into bioavailable metabolites and inorganic nutrients. Their extensive respiratory capacity and nutrient acquisition systems support N, P, and micronutrient mobilization while maintaining redox cycling. Intermediate-layer taxa may facilitate fermentative processing and cross-feeding by generating short-chain fatty acids and organic acids under fluctuating oxygen availability. Despite reduced metabolic breadth, lower-layer taxa tend to occupy specialized niches that contribute to substrate turnover, stress resilience, and effective habitat/root colonization. Together, this multi-layered MAGs-defined metabolic organization highlights a cooperative and functionally integrated microbiome in which primary degraders, fermentative intermediates, and specialized taxa collectively drive redox-linked nutrient turnover, metabolic cross-feeding, and plant growth–promoting interactions that could be translated through the indigenous Jeevamrit formulation application in cost-effective, sustainable agriculture.

## Materials and Methods

### Jeevamrit metagenome assembly and recovery of MAGs

Community metagenomic DNA from Jeevamrit, prepared under different mixing regimes (constant mixing and no mixing; here, mixing served as a proxy for oxygenation extent), was extracted and sequenced on an Illumina NovaSeq 6000 using 2x250 bp paired-end chemistry (300 cycles). The demultiplexed reads were processed within the SqueezeMeta v1.3.0 pipeline (62), which integrates Trimmomatic v0.36 for adapter and primer removal, quality trimming (Phred <25), and filtering of short or ambiguous reads, followed by assembly with MEGAHIT, binning using CONCOCT v1.1.0 and MetaBAT2 v2.12.1, and bin refinement with DAS Tool to obtain optimized metagenome-assembled genomes (MAGs). Quality assessment was performed using QUAST, while MAG completeness and contamination were evaluated using CheckM, both implemented via the Galaxy platform (https://usegalaxy.org). MAGs with ≥90% completeness and ≤15% contamination were selected for downstream analyses.

### Taxono-genomic and comparative functional analyses of MAGs providing a nexus to PGPB/R traits

Taxonomic profiling of CM and NM metagenomes was performed using Kaiju against the NCBI-nr protein database, with assignments summarized across taxonomic ranks for comparison, as implemented in the KBase platform (https://kbase.us). Near-full-length 16S rRNA gene sequences of each MAG were searched for homology against validly published taxa using EZBioCloud (https://www.ezbiocloud.net/) to demarcate taxonomically closest relatives. Following functional annotation with RASTtk (using SEED-based subsystems; https://rast.nmpdr.org/), single-copy marker gene (*gyrB, rpoB, recA*) sequences were extracted and searched for homology using NCBI BLASTp, as these genes provide higher taxonomic resolution by avoiding copy-number bias relative to highly conserved markers (63). Maximum-likelihood-based phylogenetic reconstructions were performed using MEGA v11 (64) with 1,000 bootstrap replicates. The relative distribution of taxa corresponding to the MAGs was analyzed (in CM and NM communities) and visualized using the tvBOT module in ChiPlot (65).

Genome-guided taxonomic placements (average nucleotide identity and *in-silico* DNA-DNA hybridization) of the MAGs were performed using various indices. Average nucleotide identity (ANI) was assessed by OAT (66), which estimates pairwise nucleotide similarity based on orthologous genomic fragments. Type (Strain) Genome Server (TYGS) (https://tygs.dsmz.de/) based typing was performed for genome-based taxonomic placement against type strain genomes and derivation of DNA-DNA hybridization (dDDH) values . To further validate its standardized genomic relatedness and correct phylogenomic placement, genome-based taxonomic demarcation was performed using GTDB-Tk based on reference genome associations, as implemented in the KBase platform (https://kbase.us).Circular genome maps (with details of size, strand orientation, organization, gene transcription extent, GC skew, and resistance gene markers) were generated using CGView in the Proksee server (https://proksee.ca/), whereas Fast-ANI was used to assess genomic synteny rearrangements.

Functional mapping of MAGs gene categories was performed using eggNOG-mapper in OmicsBox v3.0.30 (https://www.biobam.com/omicsbox/). Functional differences among genome size–normalized COG categories were evaluated using a Friedman test with MAG as a blocking factor. Effect size was quantified using Kendall’s W followed by paired Wilcoxon tests with Benjamini-Hochberg correction in R (v2024.12.1). The metabolic potential of recovered MAGs was further predicted using the DRAM v1.4.5 tool in KBase (https://www.kbase.us/), considering core metabolic pathways, redox processes, and ecological functions. Biosynthetic gene clusters were identified using antiSMASH v7.0 (https://antismash.secondarymetabolites.org/#!/start) to predict secondary metabolite biosynthetic potential. Carbohydrate-active enzymes were mined against the CAZy database (db-CAN) (https://pro.unl.edu/dbCAN2/) deployed in the KBase. Genes responsible for exerting plant growth– promoting (PGP) traits were identified from SEED subsystem annotations and using the open-source PLaBase-PGPT-Pred platform (https://plabase.cs.uni-tuebingen.de/). Gene abundances across major metabolic pathways were quantified and visualized as heatmaps using the ComplexHeatmap package in R (v4.3.1).

## Data availability

Metagenomic data are available in the NCBI Sequence Read Archive under accession numbers SRR33746293 and SRR33746294. The MAGs were submitted under the bioproject PRJNA1269043 and biosample (SAMN54954738 to SAMN54954753).

## Acknowledgments

The authors acknowledge Gujarat Biotechnology University for providing the necessary research facilities. This study was funded by the Gujarat State Biotechnology Mission (GSBTM), under the Network Research Program on Natural Farming through Biotechnology Interventions by the Department of Science and Technology, Government of Gujarat, India (Grant ID: GSBTM/JD(R&D)/661/2022-23/00173054).

The authors declare no conflicts of interest.

